# PastView: a user-friendly interface to explore evolutionary scenarios

**DOI:** 10.1101/651661

**Authors:** François Chevenet, Guillaume Castel, Emmanuelle Jousselin, Olivier Gascuel

**Affiliations:** MIVEGEC, Université de Montpellier, CNRS, IRD, France; LIRMM, Université de Montpellier, CNRS, France; CBGP, INRA, CIRAD, IRD, Montpellier SupAgro, Université de Montpellier, France; Unité de Bioinformatique Evolutive, C3BI, USR 3756, Institut Pasteur, CNRS, Paris, France

**Keywords:** phylogeny, ancestral character states, evolutionary scenario

## Abstract

**Background:** Ancestral character states computed from the combination of phylogenetic trees with extrinsic traits are used to decipher evolutionary scenarios in various research fields such as phylogeography, epidemiology, and ecology. Despite the existence of powerful methods and software in ancestral character state inference, difficulties may arise when interpreting the outputs of such inferences. The growing complexity of data (trees, annotations), the diversity of optimization criteria for computing trees and ancestral character states, the combinatorial explosion of potential evolutionary scenarios if some ancestral characters states do not stand out clearly from others, requires the design of new methods that operate on tree topologies and extrinsic traits associations to ease the identification of evolutionary scenarios.

**Result:** We developed a tool, PastView, a user-friendly interface that includes numerical and graphical features to help users to import and/or compute ancestral character states and extract evolutionary scenario as a set of successive transitions of ancestral character states from the tree root to its leaves. PastView offers synthetic views such as transition maps and integrates comparative analysis methods to highlight agreements or discrepancies between methods of ancestral annotations inference.

**Conclusion:** The main contribution of PastView is to assemble known numerical and graphical methods into a multi-maps graphical user interface dedicated to the computing, searching and viewing of evolutionary scenarios based on phylogenetic trees and ancestral character states. PastView is available publicly as a standalone software on www.pastview.org.

## Background

Phylogenetic trees are often combined with extrinsic traits to reconstruct evolutionary scenarios in various research fields, including phylogeography, molecular epidemiology, and ecology. The analysis starts with the computation of ancestral annotations from extrinsic traits (discrete variables such as geographic origin, resistance to a treatment, and life history traits), associated with the sampled sequences used to build the phylogenetic tree. It continues with the study of the evolutionary changes from the tree root to its leaves to characterize evolutionary scenarios such as the spread of a disease, the dynamic of a drug resistance, or shifts in ecological habitats.

There are numerous methods and software applications in ancestral character state inferences, among the most used are SIMMAP [1], BEAST [2], SpreaD3 [3], and various functions implemented in R packages [4, 5]. However, many difficulties may arise when interpreting the outputs of these methods. First, there is a diversity of optimization criteria for computing trees and ancestral annotations (parsimony, maximum likelihood, Bayesian inference) and each of them also involves various models of evolution; these methods may all yield different results that need to be compared. Second, the tree size and the complexity of the annotations can be an inconvenience with respect to computation time and they often require tedious graphical analyses in order to be interpreted. Third, although probabilistic models give more accurate results than other methods, if some ancestral characters do not stand out clearly from others (*i.e. if* they are not much more likely), they may produce a combinatorial explosion of potential evolutionary scenarios.

To the best of our knowledge, there are no tools available to compare different sets of ancestral annotations and to find common patterns across multiple evolutionary scenarios from a beam of ancestral annotations transitions. We thus present PastView that provides numerical and graphical tools to facilitate the interpretation of evolutionary scenarios derived from phylogenetic trees with ancestral character states.

## Implementation

PastView is written in Tcl/Tk. It is an open source, cross-platform, standalone editor available for Windows and Unix-like systems including OSX. The PastView process includes five steps (Fig. 1): (1) data input (tree and annotations), (2) tree and/or annotations edition, (3) computing and displaying of ancestral annotations, (4) study of ancestral annotations transitions and (5) comparative analysis between different methods of ancestral annotation inference. The PastView user interface/architecture (Fig. 2) of the stand-alone module is based on multi-maps, where each map is subdivided into a main view for the tree display and secondary view(s). PastView’s features are grouped into five toolboxes, according to the solving process:

1. The “Input/output” toolbox (Fig. 2c1) includes controls for I/O management: loading a phylogenetic tree (Newick format) and ancestral annotations or primary annotations (i.e. annotations for leaves only, see below for computing ancestral annotations) in CSV format. The user can also import data in the NEXUS format, which merges tree topology with ancestral annotations (*e.g.* BEAST output). PastView’s graphical output format is PostScript and/or SVG for high quality graphics and CSV for annotations.
2. The “Editing trees and annotations” toolbox (Fig. 2c2) includes controls for basic editing of trees and annotations: tree layouts, rooting, swapping, ladderizing, zooming in/out, etc.
3. The third toolbox (Fig. 2c3) is related to the “Computation and display of ancestral annotations”. PastView computes ancestral annotations using parsimony: DELTRAN and ACCTRAN [6, 7, 8] and maximum likelihood with an optimized scaling factor: F81-like marginal posteriors and joint inference [9, 10]. Ancestral annotations are displayed with color-coded bubbles and pie charts with the use of thresholds/filters to simplify views (*e.g.* highlighting contentious nodes with several annotations having high but closed probabilities, assemble annotations with small probabilities, etc.). Ancestral annotation can also be displayed with a tree foreground color-coded under the constraints of a probability threshold and/or a background color-coded according to the Size criterion (Sz, i.e. number of descendant with the same annotation, as in the PhyloType method [11]).
4. Toolbox for analysis of ancestral annotation transitions (Fig. 2c4): a transition is a change in ancestral annotation between subsequent nodes of a rooted phylogenic tree (Top-Down reading). PastView includes numerical and graphical methods to study transition suites. A summary view of all transitions observed in the tree is called a transition map (Fig. 3). There are three kinds of transition maps included in PastView. With the transition map type 1, a node is created in the transition map if a transition occurs within the phylogenetic tree. The second kind of transition map brings together similar transitions occurring at different positions and depths in the phylogenetic tree when they have the same ancestor in the transition map of type 1. For instance, in the example Fig. 3, the transition from « gray square » to « red circle » is observed twice in layout 1 (nodes t1 and t2), they are then collapsed into one node in layout 2. Finally, the transition map type 3 brings together identical transitions occurring at different positions and depths in the phylogenetic tree when they have the same ancestor in the transition map of type 2. Transitions maps can be displayed with different layouts (rectangular, slanted, and radial) and can be drawn dynamically to show the transitions depending on the node distance to the tree root (transition map type 1 only). Conversely, pointing at a node on a transition map automatically highlights the corresponding part(s) of the phylogenetic tree. The transition maps can be saved in Newick format. Transition maps are also available with a web interface on www.pastview.org. The transition matrix provides another overview of transitions, crossing annotations and giving a count or a more elaborated index of transitions between pairs of annotations (for instance, see [12]). PastView includes two types of matrices. The first one computes the number of transitions in the tree, from one ancestral annotation to another. The second one is a fast, count-based estimation of the relative transition rates, where raw counts are normalized and divided by state priors. PastView offers a query system to search for a given evolutionary path: the user selects a transition sequence (wild cards may be used) and matching tree pathways are identified and highlighted. For example, the scenario “A * B” lights up the tree pathways entailing the transitions from “A” to “B”, independently of the intermediaries (example in Fig. 4e).
5. Comparative analyses: this toolbox (Fig. 2c5) is dedicated to comparing multiple datasets (trees/ancestral annotations). For example, given a tree and ancestral annotations resulting from different inferences, PastView automatically adjusts tree foreground colors for nodes with similar results, but highlights nodes (bubbles, pie charts) exhibiting incongruences in ancestral annotations (example in Fig. 5a).

**Figure 1.**
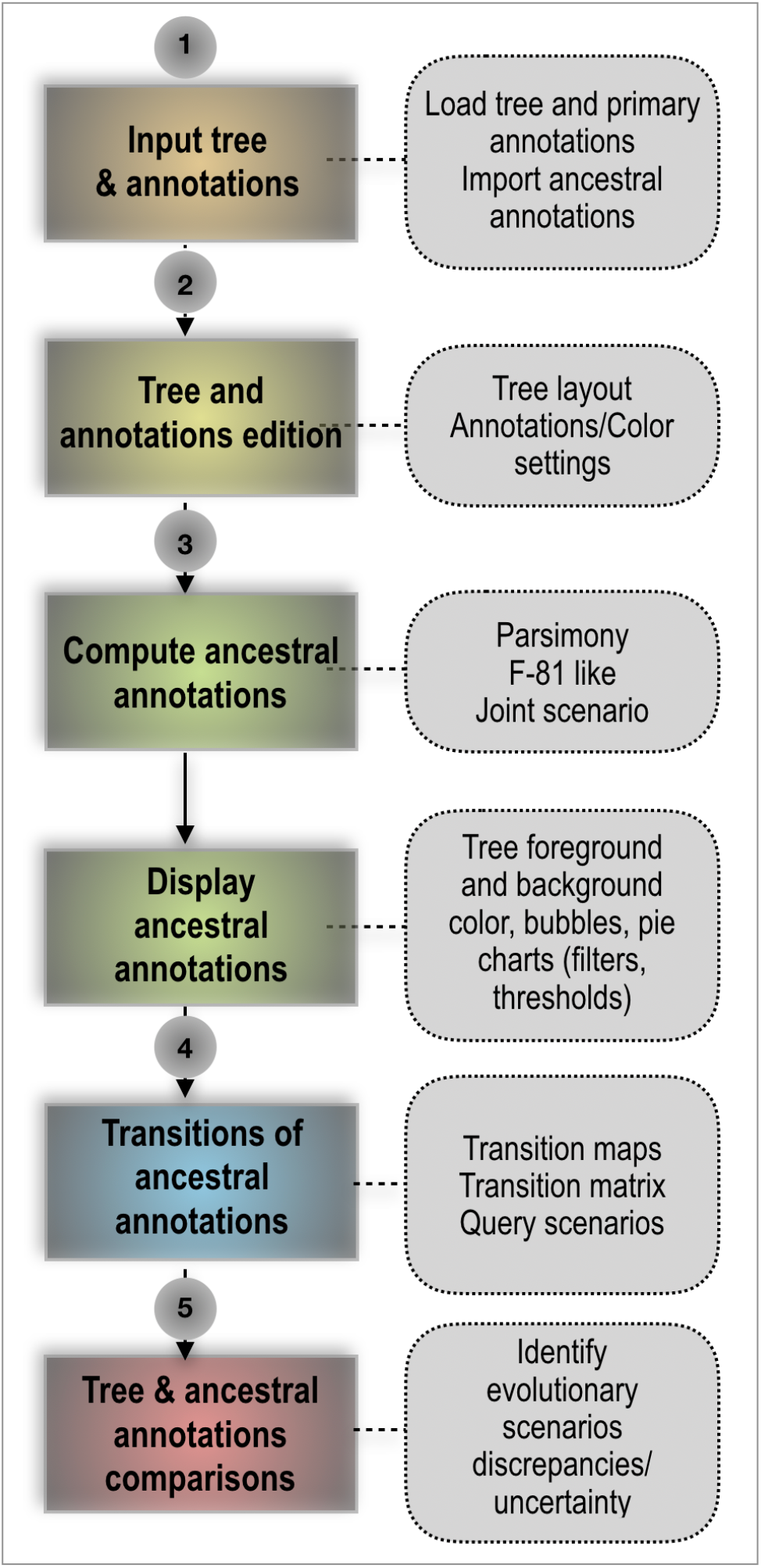
PastView solving process. The solving process of PastView includes five steps (1-5). The first step is dedicated to the input of an analysis (*e.g.* loading a tree and its annotations, importation of ancestral annotation). Step 2 is related to tree and annotations edition (settings of colors to annotations, tree view management, etc.). The step 3 of the solving process is related to the computing and displaying of ancestral annotations. Step 4 is dedicated to the study of ancestral annotations transitions. The last step is related to comparative analysises, such as visualizing different ancestral annotations sets for a given tree.

**Figure 2.**
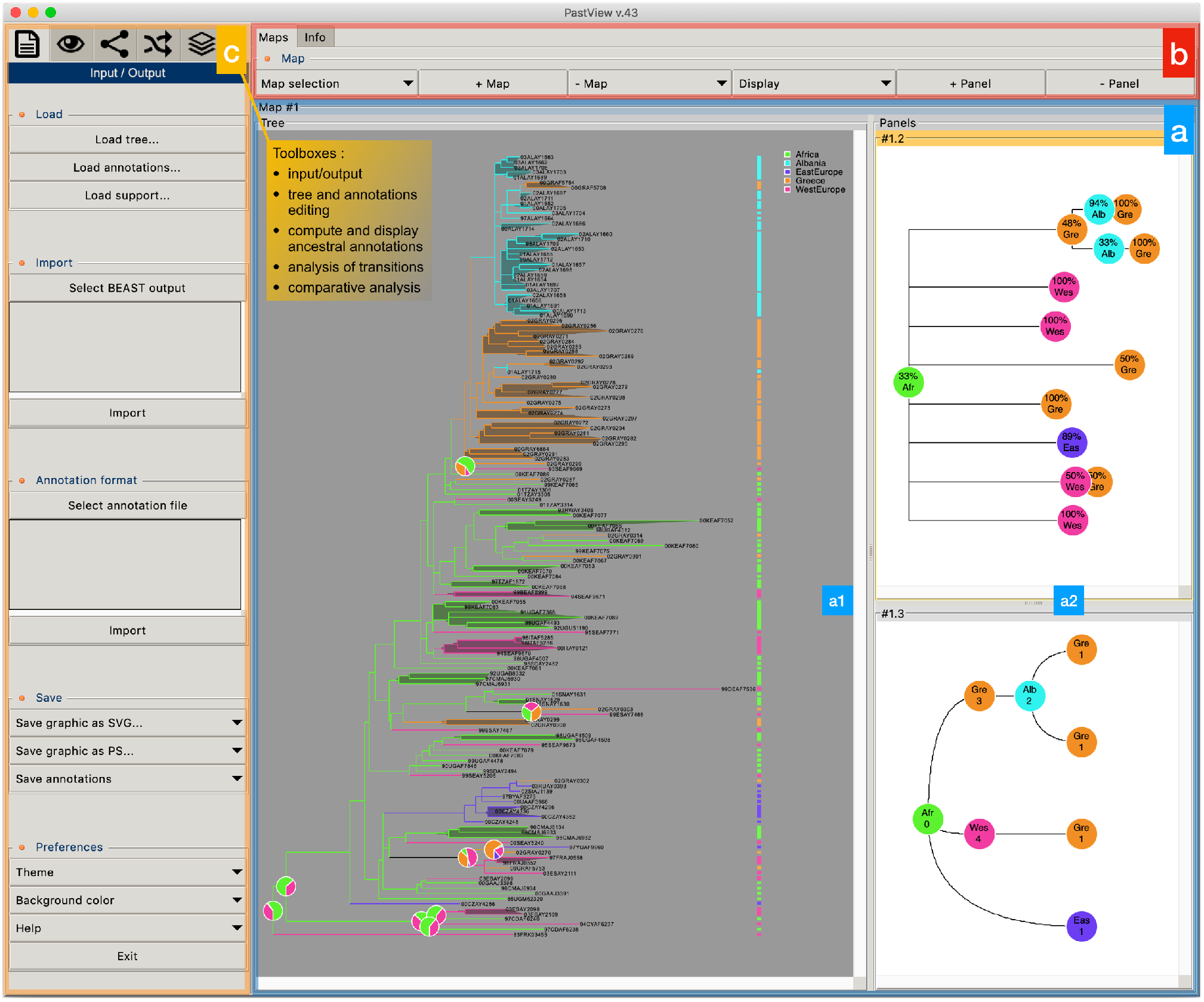
PastView user’s interface, « stand-alone » module. **(a)** the PastView interface is composed of multiple maps, where each map is subdivided into **(a1)** a main « tree view » and **(a2)**, one or several secondary panels. **(b)** the controls for the management of maps and panels. **(c)** the controls for the PastView analysis are gathered into five toolboxes, following the five steps of the problem solving process : input/output (loading files, output graphics), tree and annotation editing (annotation color settings, tree rooting, swapping, etc.), compute and display ancestral annotations, analyses of transitions and comparative analyses.

**Figure 3.**
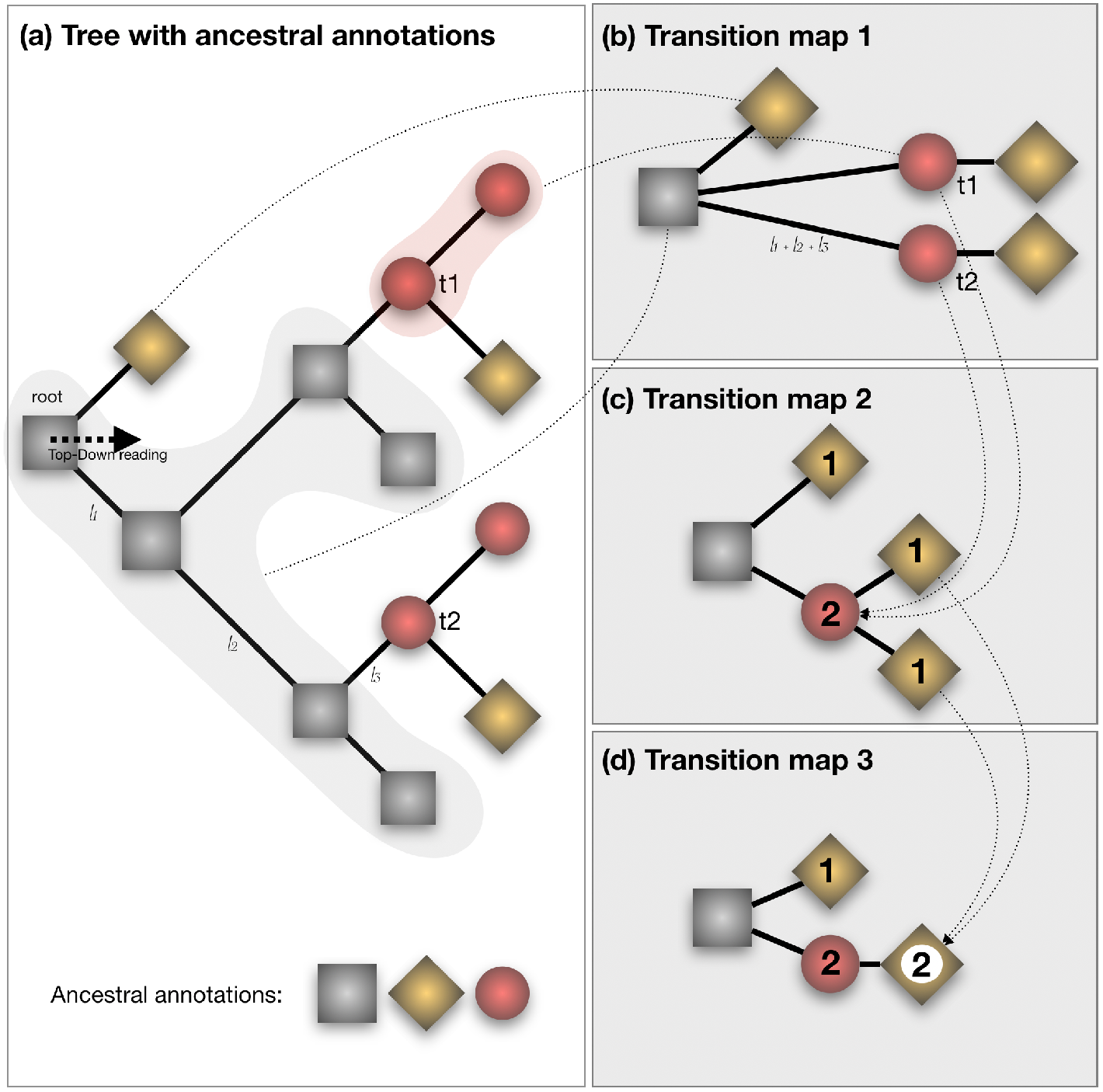
Tree-like representations of transitions. **(a)** phylogenetic tree with ancestral annotations (gray squares, red circles and gold diamonds). **(b),(c)** and **(d)**: corresponding transition maps. A transition is a change of an ancestral annotation along a phylogenetic tree (from the root to the leaves). The transition map layout 1 **(b)** is a multifurcating tree-like representation of transitions. A node is created in the map foreach transition observed in the phylogenetic tree (with the exception of the root). It can take into account branch lengths of the tree (sum of branch lengths along the path). The transition map layout 2 **(c)** gather transitions when they share the same father node in the layout 1. The number of transitions collapsed is then displayed (1 from « gray square » or « red circle » to « gold diamond », 2 from « gray square » to « red circle »). The same collapsing process is applied in turn to layout 2 to produce the transition map layout 3. Transition maps can be computed for different sets of ancestral annotations (*e.g.* different inference methods). When the output consist into probability distributions (*e.g.* Bayesian or F81-like methods), only the annotation having the greatest probability value is considered.

**Figure 4.**
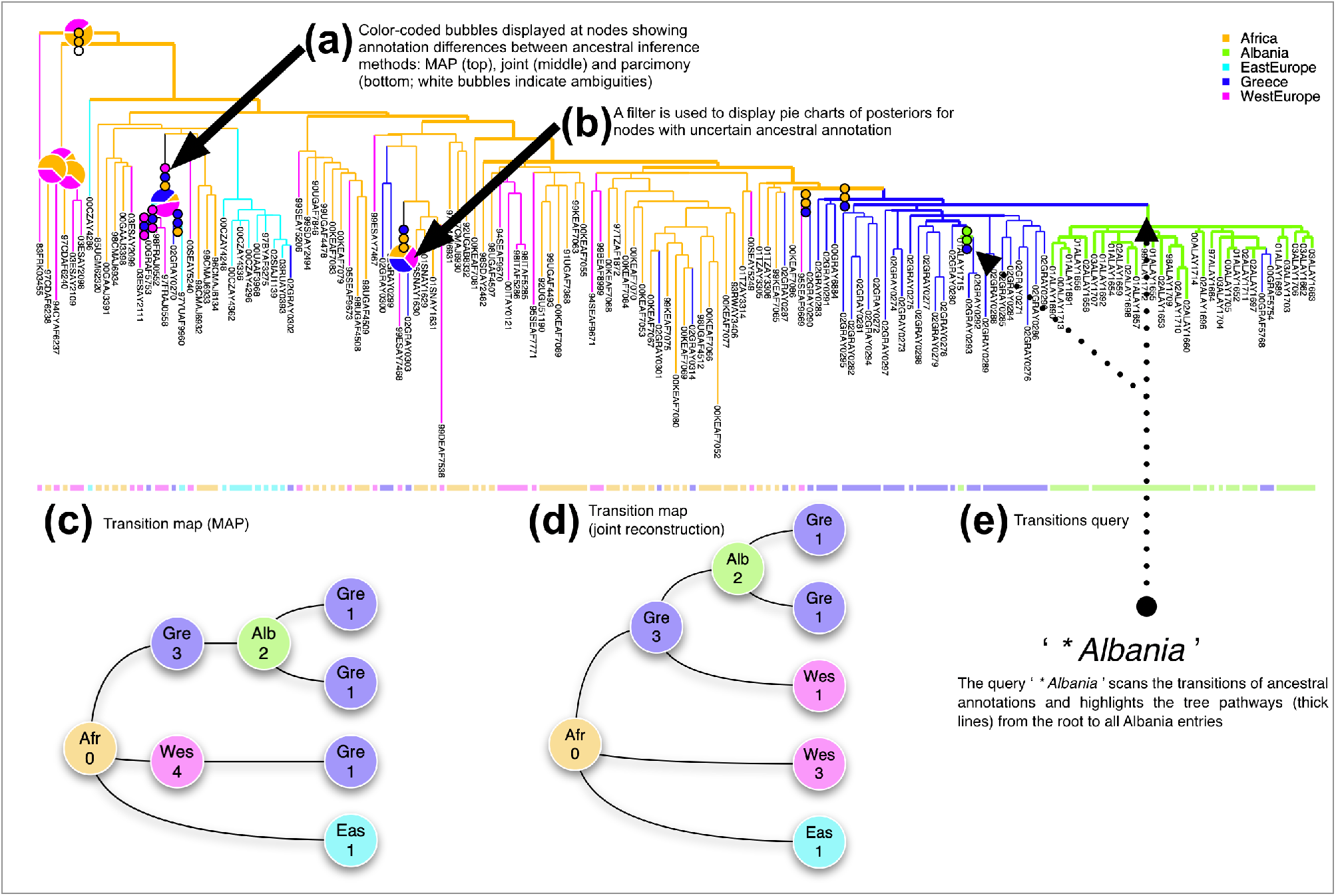
Output of a PastView analysis for the study of HIV-1A epidemiological history in Albania (Salemi *et al.*, 2008). The countries associated with the tree sequences are used to compute ancestral areas by three methods: maximum a posteriori (MAP), the joint most likely scenario, and parsimony (DELTRAN). (a) A country color-code is used to color nodes and branches if their associated ancestral annotations are the same for the three methods; if not, bubbles are displayed. (b) A filter (threshold based on MAP probability minus 40% of its value) is used to display pie charts of posteriors for ambiguous nodes. (c), (d) Tree-like representations of transitions (MAP and joint inferences, respectively); numbers indicate the numbers of identical transitions having the same ancestor in the transition maps type 1. (e) The transition query ‘* Albania’ highlights the tree pathways from the root to Albania; it displays the two distinct transitions from Greece to Albania.

**Figure 5.**
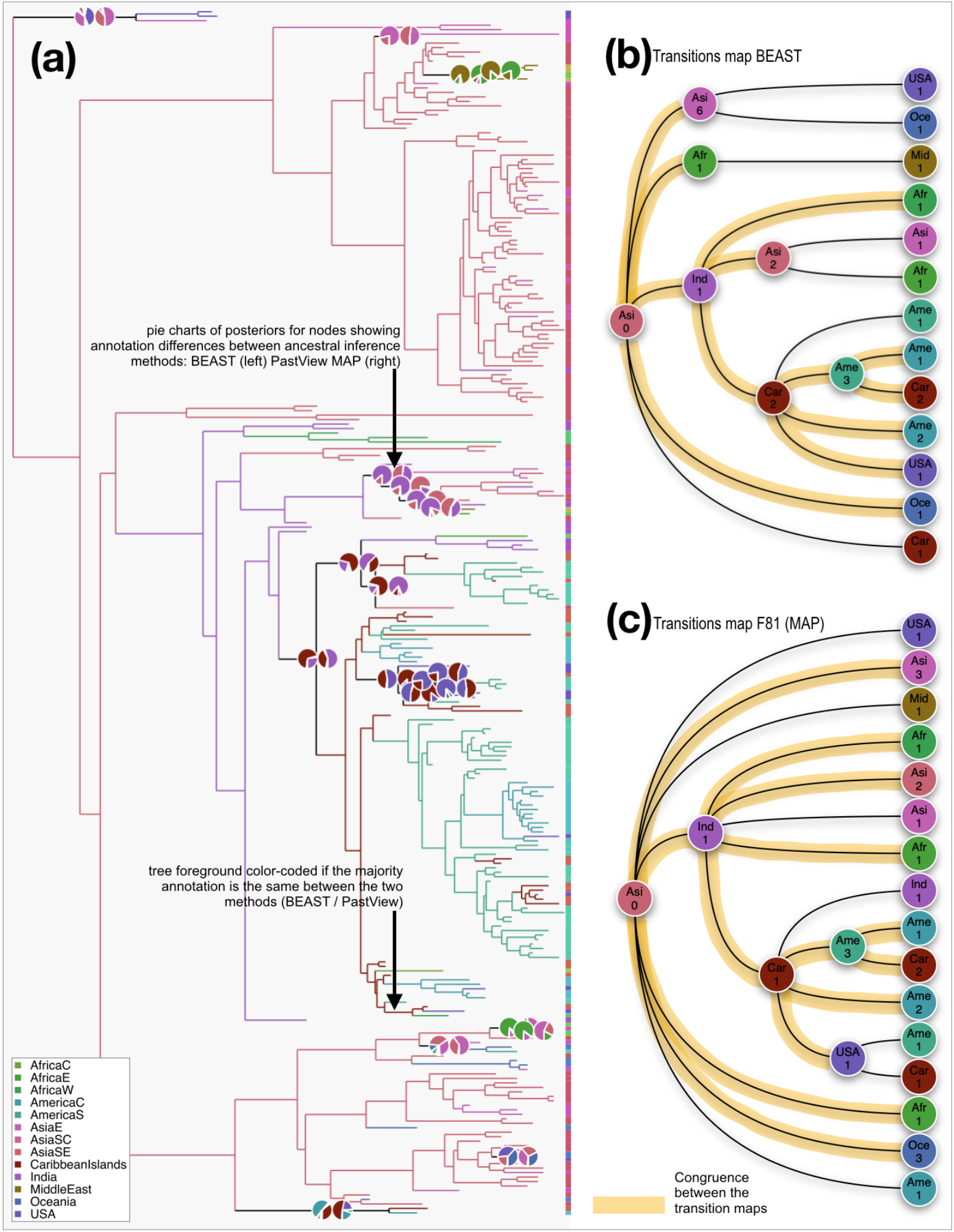
Output of a PastView analysis for the study of the global phylogeography of Dengue type 1 virus. The tree with its ancestral locations is imported from a BEAST output computed from the dataset used in (Walimbe *et al.*, 2014). Ancestral locations are then also computed with PastView maximum a posteriori method (MAP). (a) a country color-code is used to color the tree when the associated ancestral annotations (majority) are the same between BEAST and MAP (F81-like); if not, pie charts are displayed (root to tip reading: BEAST then PastView MAP). (b and c) the transitions maps are computed for each method. Compatibility between each map is highlighted (orange foreground color).

## Results

In the following, we present two published examples of virus phylogeographic studies re-examined with PastView. The first one is related to the HIV-1A epidemiological history in Albania. The second is related to the global phylogeography of Dengue type 1 virus.

### Exemple 1. Study of HIV-1A epidemiological history in Albania

Figure 4 shows a simple example of PastView output, in the study of HIV-1A epidemiological history in Albania. The dataset, tree, and origin of the sequences (locations) are from [13] (see also [11]). Ancestral locations are computed by parsimony (DELTRAN), maximum of the marginal posterior probabilities (MAP, F-81 like), and joint most likely scenario. Computation time of ancestral annotations for this small dataset (153 strains) is ~2s with a 3.1 GHz Intel I7 computer. The tree is foreground color-coded if the ancestral annotations are identical between MAP F-81 like, joint most likely reconstruction and parsimony. If not, color-coded bubbles are displayed (Fig. 4a) (in the same order, reading from root to tips). Here, we identify nine nodes with discrepancies, and only four nodes if we just compare only MAP and joint inferences. The pie charts display (Fig. 4b) the posteriors for nodes having two or more annotations with a probability higher than their MAP F-81 like probability, minus 40% of their value. Based on MAP and joint inferences, the transition maps (Fig. 4c and 4d) are slightly different but output the same global scenario: the virus spread from Africa to West Europe, East Europe and Greece, and then from Greece to Albania, with a few Greek sequences coming back from Albania. This reaches the same general conclusion as [13]: there has been a single major introduction of HIV-1A from Greece followed by a local epidemic spread. This result (Fig. 4e) is highlighted by the thick pathways from the tree root to all entries to Albania.

### Exemple 2. Study of the global phylogeography of Dengue type 1 virus

The second example is related to the study of the worldwide phylogeography of Dengue virus serotype 1 (DENV-1). The dataset, tree (269 strains), and origin of the sequences (13 locations) are the same than in from [14]. Tree and ancestral annotations are first computed with BEAST (same parameters / model as described by [14]), then imported with PastView. Another set of ancestral annotations is computed with PastView (F81-like method, MAP, computation time is ~40s with an 3.1 GHz Intel I7 computer). The tree is then foreground color-coded (Fig. 5.a) if the ancestral annotations are the same between the two methods. If not, pie-charts are displayed (from a root to tips reading: left= BEAST, right=PastView). Both analyses reach the same main conclusions as described in [14]: Southeast Asia countries are found to be the most likely origin of the virus dispersion and India played a crucial role in the establishment, evolution and dispersal of the Cosmopolitan (Africa, America, Carribean, East & Southeast Asia) DENV-1 genotype (Fig. 5b and 5c). The Caribbean region is also found by both methods as the dissemination origin point of the virus to the New world countries (South and Central America, then North America). If there is a global consensus for the most ancestral nodes/ transitions, some differences between the results obtained with the two methods exist especially for the more recents nodes/transitions (Fig. 5a, 5b & 5c). These disagreements are related mainly to transitions within small clusters of sequences annotated in heterogeneous ways.

## Conclusion

The main contribution of PastView is to assemble known numerical and graphical methods/tools into a multi-map graphical user interface dedicated to computing, searching and viewing evolutionary scenarios using phylogenetic trees and ancestral character states. One challenge will be now to develop and integrate new methods to compute ancestral annotations obtained from the nuanced outputs of the ML/Bayesian methods and the overly stringent output of the joint method, such as the recent PastML [15]. In the long-term, a next step will be to integrate statistical tools and information visualization methods to quickly identify robust evolutionary scenarios, such as a consensus of transition maps between different methods (tree computations, ancestral annotation inferences), or for more elaborated transition maps in the case of huge datasets. In the context of a growing complexity of data (*e.g.* massive data from high throughout sequencing low cost sequencing), it is easy to imagine situations where trees with tens of thousands of sequences (*e.g.* HIV), and complex annotations in quantity and quality (*e.g.* “Gene Ontology », any data related to patients), are used and without evidence of a strong structuring of data between tree and annotations. It becomes thus necessary to design methods that operate on tree topologies / extrinsic traits associations and to produce synthetic views, to extract evolutionary scenarios.

## Availability of data and materials

**Project name**: PastView

**Project home page**: http://pastview.org/

**Operating system(s)**: Platform independent

**Programming language**: Tcl/Tk

**Other requirements**: ActiveTcl 8.6.8 or higher

**License**: GNU GPL

**Any restrictions to use by non-academics**: none

## Availability of data and materials

PastView source code and data (tree, primary and ancestral annotations) of the two examples used in the manuscript are available on http://pastview.org/ and in the PastView package.

## Acknowledgements

We would like to thank S. Guindon for providing insightful ideas on this topic.

## Availability of data and materials

PastView and the source code is publicly available on www.pastview.org

## Authors’ contributions

FC conceived the idea, developed the tool and wrote the manuscript.

OG conceived the idea, participated in supervision of the project.

EJ participated in the idea and tested the tool

GC participated in the idea and in supervision of the project and tested the tool

All authors have read, revised, and approved the final manuscript.

## Ethics approval and consent to participate

Not applicable

## Consent for publication

Not applicable

